# Decreasing pdzd8-mediated mitochondrial-ER contacts in neurons improves fitness by increasing mitophagy

**DOI:** 10.1101/2020.11.14.382861

**Authors:** Victoria L. Hewitt, Leonor Miller-Fleming, Simonetta Andreazza, Francesca Mattedi, Julien Prudent, Franck Polleux, Alessio Vagnoni, Alexander J. Whitworth

## Abstract

The complex cellular architecture of neurons combined with their longevity makes maintaining a healthy mitochondrial network particularly important and challenging. One of the many roles of mitochondrial-ER contact sites (MERCs) is to mediate mitochondrial quality control through regulating mitochondrial turn over. Pdzd8 is a newly discovered MERC protein, the organismal functions of which have not yet been explored. Here we identify and provide the first functional characterization of the *Drosophila melanogaster* ortholog of Pdzd8. We find that reducing pdzd8-mediated MERCs in neurons slows age-associated decline in locomotor activity and increases lifespan in *Drosophila*. The protective effects of pdzd8 knockdown in neurons correlate with an increase in mitophagy, suggesting that increased mitochondrial turnover may support healthy aging of neurons. In contrast, increasing MERCs by expressing a constitutive, synthetic ER-mitochondria tether disrupts mitochondrial transport and synapse formation, accelerates age-related decline in locomotion and reduces lifespan. We also show that depletion of pdzd8 rescues the locomotor defects characterizing an Alzheimer’s disease (AD) fly model over-expressing Amyloidβ_1–42_ (Aβ_42_) and prolongs the survival of flies fed with mitochondrial toxins. Together, our results provide the first *in vivo* evidence that MERCs mediated by the tethering protein pdzd8 play a critical role in the regulation of mitochondrial quality control and neuronal homeostasis.

## Introduction

Since the vast majority of neurons are postmitotic, maintaining functional neurons throughout an organism’s lifetime requires tight regulation of organelle functions and stress responses. Mitochondria and the endoplasmic reticulum (ER) extend throughout neuronal processes including axons and dendrites, and both are vital and interdependent contributors to neuronal health (Wu *et al*, 2017). Mitochondria-ER contacts (MERCs) are controlled by a variety of contact site proteins and contribute to a range of functions required for proper development and maintenance of postmitotic neurons, including regulation of calcium homeostasis, lipid biogenesis, organelle reshaping and dynamics, and metabolic signalling (Giacomello & Pellegrini, 2016; Paillusson *et al*, 2016). Dysregulation of MERCs is particularly damaging to neurons as they are particularly susceptible to calcium overload, oxidative and ER stresses and to altered mitochondrial function, localization and transport (Lee *et al*, 2018a; Misgeld & Schwarz, 2017).

As MERCs are modulated by a number of different protein complexes, the detrimental effects of dysregulation of MERCs are varied due to the diversity of these contact site functions (Martino Adami *et al*, 2019). The critical function of MERCs in regulating cellular responses to damage and stress is underscored by the finding that many human patient cellular models and animal models of age-related neurodegenerative diseases have been shown to have disrupted MERCs. Both reduced MERCs (De Vos & Hafezparast; Sepulveda-Falla *et al*, 2014) and increased MERCs (Area-Gomez *et al*, 2012; Gómez-Suaga *et al*, 2019; Zampese *et al*, 2011) have been implicated in neurodegenerative diseases. Consequently, there is still little consensus on how altered MERCs contribute to neurodegeneration, even within a single disease model (Erpapazoglou *et al*, 2017). The various functions of MERCs make it likely that multiple mechanisms might be involved and, with an ever-expanding toolkit, we can now better define the molecular identities and specific functions of ER-mitochondria tethering complexes and begin to unify many of the seemingly conflicting discoveries in this rapidly growing field (Csordas *et al*, 2018).

Pdzd8 is one of the most recently discovered proteins that mediates mammalian MERCs (Hirabayashi *et al*, 2017) and is a paralog of Mmm1 (Wideman *et al*, 2018), a component of the fungal-specific ER mitochondria encounter structure (ERMES) and first MERC complex identified (Kornmann *et al*, 2009). We used *Drosophila melanogaster* to study the consequences of neuronal depletion of pdzd8 both at the cellular and at the organismal level. Importantly, we also describe how the phenotypes associated with neuron-specific depletion of pdzd8 change with age and may contribute to healthy aging. We show the MERCs mediated by the pdzd8 tethering protein regulate mitochondrial turnover through mitophagy and that reducing these contacts prolonged locomotor activity and lifespan in *Drosophila melanogaster*.

## Results

### Characterization of fly pdzd8

The *Drosophila melanogaster* gene *CG10362* encodes an uncharacterized protein in the PDZK8 family (Lee & Hong, 2006). *CG10362* has a similar predicted domain structure to mammalian Pdzd8 (Figure 1A). Expression of *CG10362* in flies is low but is most highly expressed in the central nervous system (FlyAtlas 2; Figure S1A)(Leader *et al*, 2018) and is enriched in neurons over glia (Figure S1B) (Davie *et al*, 2018). This specificity in expression in *Drosophila* provided an excellent opportunity to explore the neuronal functions of this newly discovered MERC protein and also to investigate the functional relevance of MERCs in adult and aging neurons. We propose that *CG10362* encodes the fly ortholog of mammalian Pdzd8 and, based on the data presented in this paper, we propose to name it *pdzd8*.

**Figure 1.**
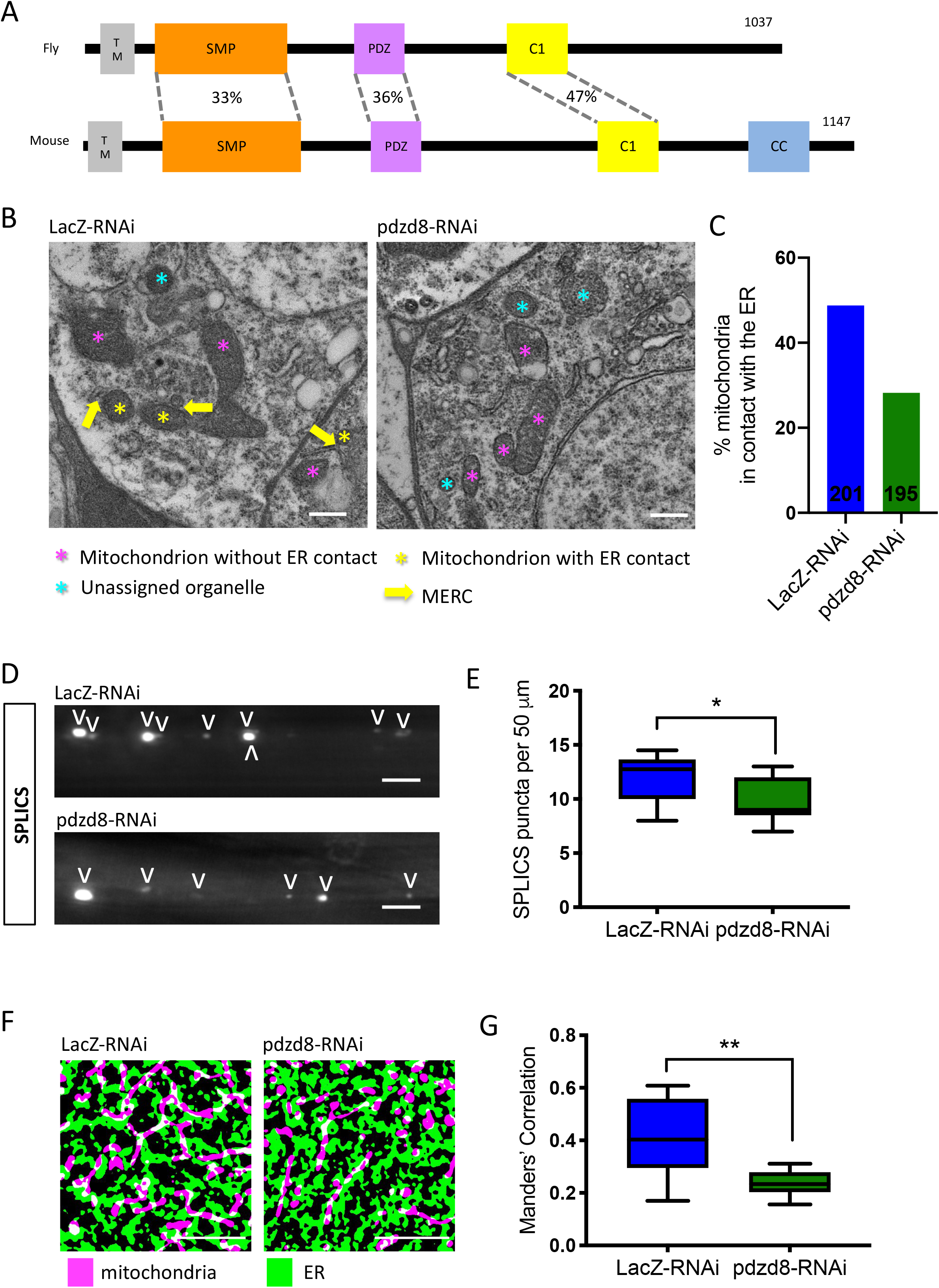
Expression of *pdzd8*-RNAi reduces mitochondria-ER contacts. (**A**) Domain organisation of Drosophila pdzd8 (CG10362) compared to mouse Pdzd8 showing percentage identities of conserved domains based on Clustal Omega alignments. Overall percentage identity of the amino acid sequences is 21 %. SMP (Synaptotagmin-like mitochondrial lipid-binding proteins) 33 % identical, PDZ (PSD95/DLG/ZO-1) 36 % identical, C1 (C1 protein kinase C conserved region 1 also known as Zn finger phorbol-ester/DAG-type signature) 47 % identical; TM: predicted transmembrane domain, CC: coil-coil domain. (**B**) Electron microscopy images of 2-day-old adult brains showing representative images of ER, mitochondria and MERCs in soma from nSyb>*LacZ*-RNAi and nSyb>*pdzd8*-RNAi flies. Scale bar 500 nm. Mitochondria without identifiable ER contacts marked with magenta *, mitochondria forming ER contact marked with yellow * with yellow arrow indicating contact location, organelles that did not contain clear cristae are marked with a cyan * and were excluded from the analysis (**C**) Percentage of mitochondria in contact with the ER from controls and panneuronal nSyb>*pdzd8*-RNAi flies quantified from EM images of 2-day-old adult brains. N = 3 brains per genotype. (**D**) SPLICS puncta indicating MERCs in axon bundles of larval neurons from controls and nSyb>*pdzd8*-RNAi flies. Quantified puncta highlighted with V. (**E**) Quantification of SPLICS puncta in C. n = 11 animals per genotype, p=0.0198, unpaired t-test with Welch’s Correction, scale bar 5 μm. (**F**) Representative binarized SIM images of ER (green) and mitochondria (purple) in larval epidermal cells from controls and da>*pdzd8*-RNAi flies. Scale bar 500 nm. (**G**) Quantification of colocalization of ER and mitochondria using Mander’s Correlations compared using an unpaired t-test with Welch’s Correction. p=0.012.

To characterize the function of pdzd8 in flies, we used the UAS-GAL4 system to manipulate its expression (Brand & Perrimon, 1993). Ubiquitous expression of an RNAi construct targeting *pdzd8* strongly reduces its mRNA levels in larvae (Figure S1C). To establish that MERCs were decreased in neurons expressing *pdzd8*-RNAi, we analysed adult fly brains by transmission electron microscopy (TEM) and manually identified contacts between ER and mitochondria (Figure 1B). Accordingly, the proportion of mitochondria in the soma of adult fly neurons in contact with ER is reduced in flies expressing *pdzd8*-RNAi (Figure 1C and S1F).

To confirm that *pdzd8*-RNAi reduces MERCs in axons as well as soma, we adapted the recently developed MERC quantification tool, a split-GFP-based contact site sensor (SPLICS) (Cieri *et al*, 2017) to create a SPLICS transgenic reporter line. The SPLICS construct targets β-strands 1-10 of GFP to the mitochondrial outer membrane and β-strand 11 to the ER membrane, and where these membranes are in close proximity fluorescent puncta are produced by reconstitution of the split-GFP (Figure S2A). To validate this tool in *Drosophila*, we expressed SPLICS in motor neurons (Figure S2B) and compared the number of puncta in control axons and those expressing a well characterized artificial ER-mitochondrial tether developed by Csordas *et al*. (Basso *et al*, 2018; Csordas *et al*, 2006). The tether construct induces formation of ~5 nm MERCs through transmembrane domains that anchor it in both the mitochondrial outer membrane and the ER (Figure S2C). The density of SPLICS puncta in the axons expressing the tether was four times higher than controls (Figure S2D), indicating the SPLICS reporter was able to detect the increased MERCs resulting from synthetic tether expression in neurons *in vivo*.

Using this SPLICS construct we also detected a significant decrease in the density of SPLICS puncta in central larval axons (bundles projecting to segments A7 and A8) expressing the *pdzd8*-RNAi compared to *LacZ*-RNAi controls (Figure 1D-E). While pdzd8 expression is low beyond the nervous system, ubiquitous knockdown of pdzd8 also reduces the extent of MERCs measured by fluorescence colocalization of ER and mitochondrial signals by super-resolution microscopy (structured illumination microscopy, SIM) in larval epidermal cells (Figure 1F-G and S1E), corroborating our results using TEM and SPLICS analysis.

Consistent with the first reported function of Pdzd8 at MERCs in mouse neurons (Hirabayashi *et al*, 2017), we observed a reduced number of MERCs (Figure 1), but no obvious changes in mitochondrial or ER morphology in fly larval or adult neurons upon pdzd8 knockdown (See Figures 1B, 1D, 1F, S1D-E, 4A-B, 4F-I, 5A and 5C). Together, these data show that *CG10362* encodes the *Drosophila* homolog of Pdzd8 and functions like mammalian Pdzd8 to regulate MERCs. We next sought to investigate the consequences of loss of pdzd8 from neurons on organismal phenotypes.

### Reduced mitochondria-ER contacts are protective in aging neurons

Knockdown of *pdzd8* in fly neurons produces viable adults. There was no impact on the locomotor performance of pan-neuronal *pdzd8*-RNAi knockdown in young flies assessed in a climbing assay (Figure 2A). Surprisingly, we found that loss of pdzd8 dramatically slowed the age-associated decline in locomotor activity assessed by the climbing assay (Figure 2A). Importantly, this effect was reproduced by motor neuron-specific *pdzd8* knockdown (Figure S3A). Strikingly, the increase in locomotor activity was accompanied by a significant increase in lifespan (Figure 2C).

**Figure 2.**
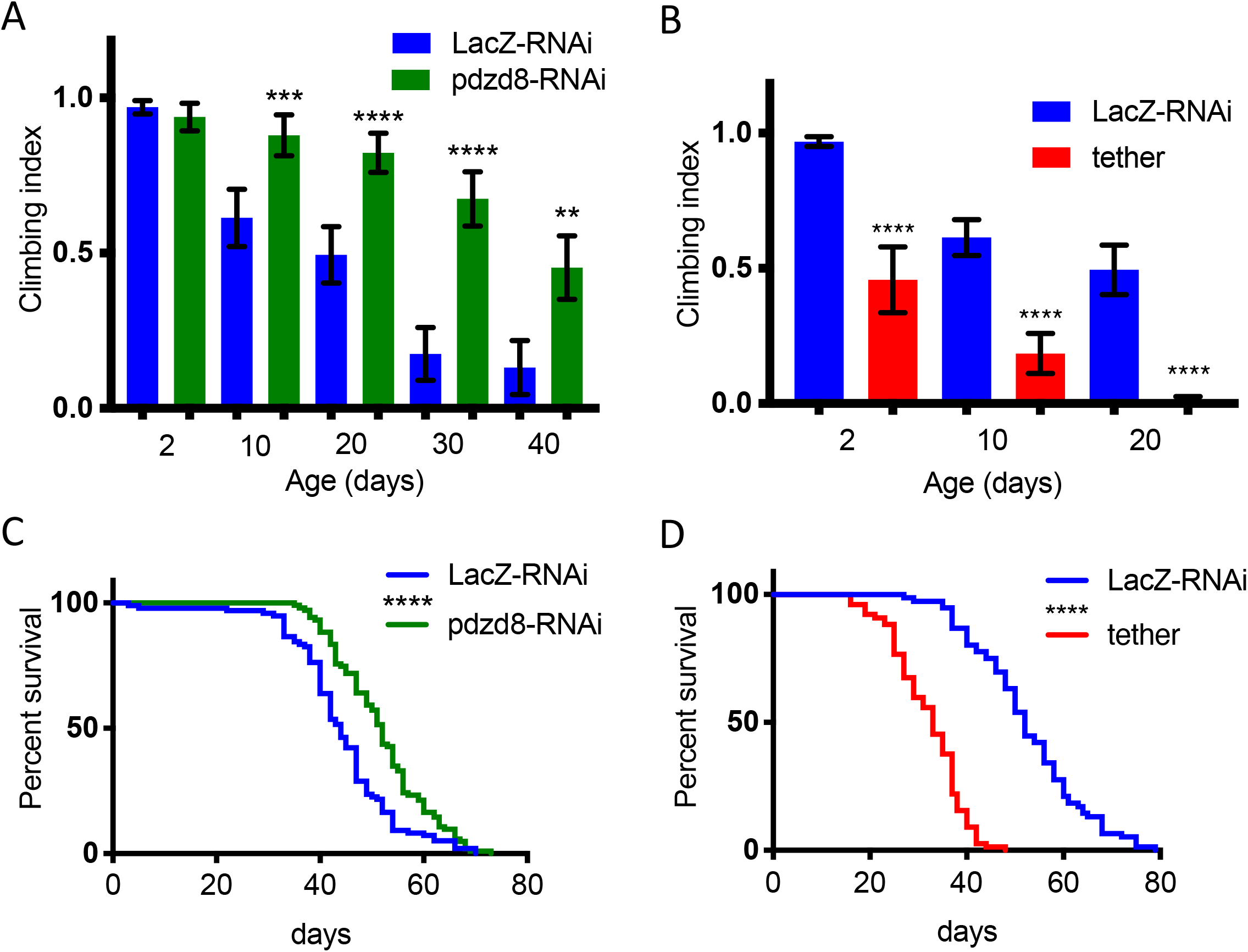
Lifespan and locomotor activity changes in aged flies with pan-neuronal (nSyb) driven alterations in tethering. (**A, B**) Locomotor activity of flies was assessed during aging by negative geotaxis climbing assays on the indicated days. n>50 flies per genotype. Flies expressing (**A**) *pdzd8*-RNAi or (**B**) synthetic tether were compared to *LacZ*-RNAi controls. (**C, D**) Lifespans in standard growth conditions and food. (**C**) Flies expressing *pdzd8*-RNAi were compared to *LacZ*-RNAi controls. n = 97, 108 per genotype, median survival 44 vs 52 days, p<0.0001. (**D**) Flies expressing the synthetic tether were compared to *LacZ*-RNAi controls. n = 74, 85 per genotype, median survival 52 vs 33 days, p<0.0001.

In contrast, increasing MERCs by expression of a synthetic mitochondria-ER tether in all neurons resulted in a climbing defect in young flies and a significant acceleration of the age-related decline in climbing (Figure 2B), consistent with previous reports (Basso *et al*, 2018). This climbing defect was exacerbated with age (Figure 2B) and associated with a substantially reduced lifespan (Figure 2D). Consistent with these results, increased expression of *pdzd8* also resulted in decreased climbing ability with age (Figure S3B). Notably, this effect was suppressed by co-expression of the *pdzd8*-RNAi, further validating the specificity of this transgene (Figure S3B). Therefore, decreasing pdzd8-mediated MERCs in neurons prolonged lifespan and protected against locomotor decline with age, while increased MERCs in neurons were found to speed up the age-related decrease in locomotor activity and decreased lifespan.

### Loss of neuronal pdzd8 promotes survival in the presence of mitochondrial toxins

To investigate how reduction of pdzd8-mediated MERCs improved fitness i.e. prevented age-related decline in locomotor activity and increased lifespan, we assessed whether reducing *pdzd8* expression in neurons may protect from additional stresses during aging. We assessed lifespan in flies aged on food with limited nutrients (5 % sucrose, 1 % agar), and found that in contrast to flies aged on a rich diet (Figure 2C), neuronal expression of *pdzd8*-RNAi no longer extended the lifespan in comparison to controls (Figure 3A). When an additional oxidative stress was introduced by adding hydrogen peroxide to the food, the flies expressing *pdzd8*-RNAi died faster than controls (Figure 3B). Therefore, in the presence of general stresses, *pdzd8*-RNAi is also not protective.

**Figure 3.**
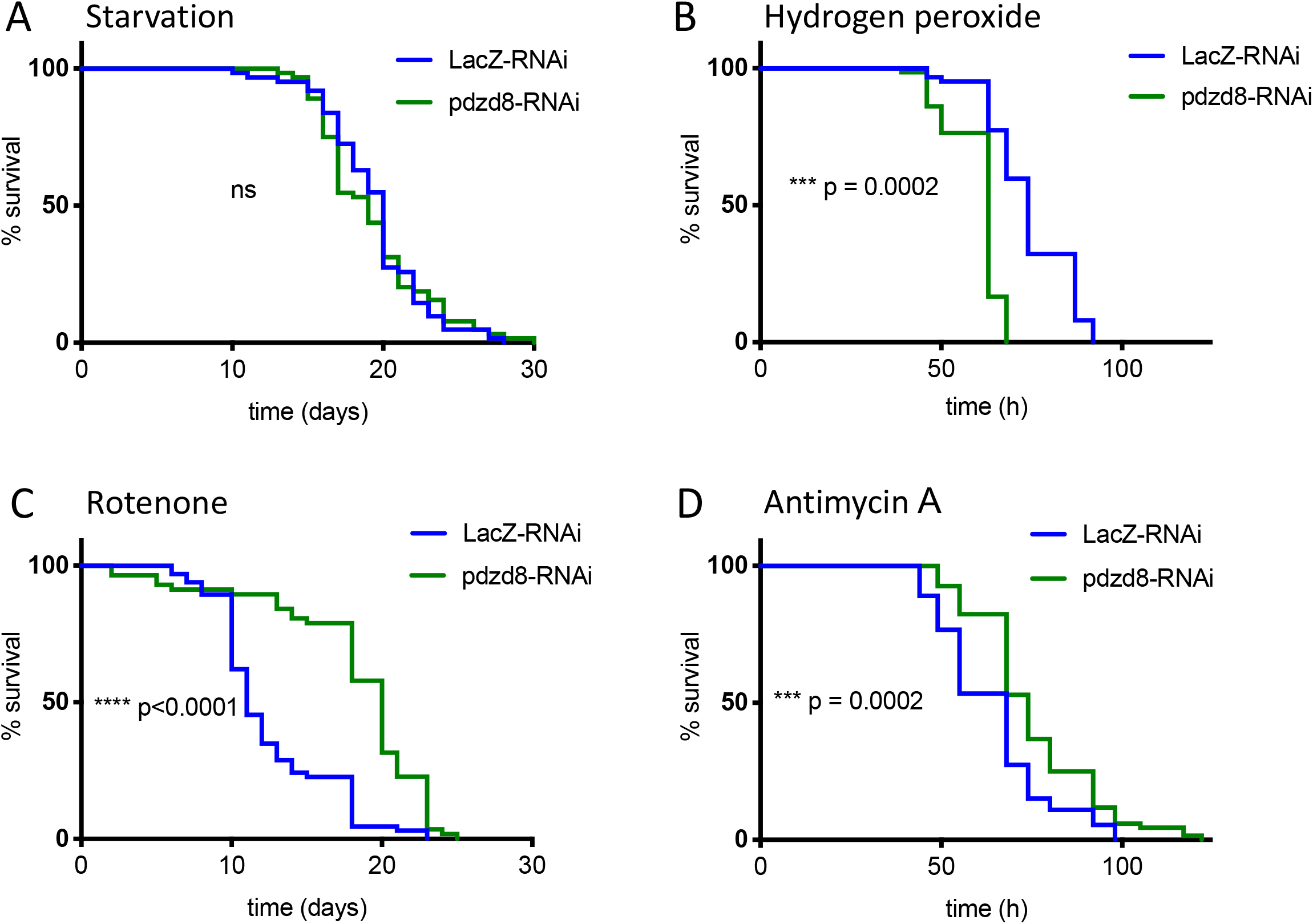
Knockdown of *pdzd8* protects flies against mitochondrial toxins. Flies expressing pan-neuronal nSyb>*pdzd8*-RNAi were compared to nSyb>*LacZ*-RNAi controls when aged on a restricted diet of food containing 1 % agar with 5 % sucrose. (**A**) Lifespan with dietary restriction alone. N = 62 vs 64, median survival 20 days vs 19 days, difference ns. (**B**) Lifespan with addition of 5 % hydrogen peroxide. Median survival: 63 h vs 74 h, n = 67, 74, p=0.0002. (**C**) Lifespan with addition of 1 mM rotenone. Median survival: 11 days vs 20 days, n = 66, 57, p<0.0001. (**D**) Lifespan with addition of 5 μg/mL antimycin A. Median survival: 74 h vs 68 h, n = 72, 68, p=0.0002.

Due to the function of pdzd8 at MERCs, we examined whether the protective effects caused by the pdzd8 depletion in neurons were associated with mitochondrial damage. To address this, we fed the flies mitochondrial toxins: Rotenone, a complex I inhibitor or antimycin A, a complex III inhibitor, both block the electron transport chain and result in dysfunctional mitochondria. Reducing pdzd8 levels in neurons significantly prolonged the survival of flies fed with mitochondrial toxins rotenone (Figure 3C) or antimycin A (Figure 3D) compared to control flies. As improved mitochondrial function could also contribute to the protective effects of loss of pdzd8 in neurons we measured ATP levels in young or aged fly heads expressing *pdzd8*-RNAi but found no significant differences (Figure S3D). These results indicated that neuronal loss of pdzd8 protects flies from damage induced by mitochondrial toxins.

### Modulating MERCs causes axonal transport and NMJ defects

While mitochondrial motility is important for neuronal health, it remains an open question whether decline of mitochondrial transport in neurons contributes to aging (reviewed in (Mattedi and Vagnoni 2019)). We first tested the hypothesis that decreased ER-mitochondrial tethering contributes to the protective effect of *pdzd8* downregulation in aging through changes in mitochondrial motility. We examined the distribution and morphology of mitochondria in axons of CCAP efferent neurons (Park *et al*, 2003). We found no significant change in mitochondrial length or density in larval axons when comparing control to *pdzd8*-RNAi or synthetic ER-mitochondria tether-expressing neurons (Figure 4A-C). However, increasing tethering dramatically decreased mitochondrial motility (Figure 4D-E). In contrast, knockdown of *pdzd8* to reduce MERCs had no effect on axonal transport in larvae (Figure 4D-E).

**Figure 4.**
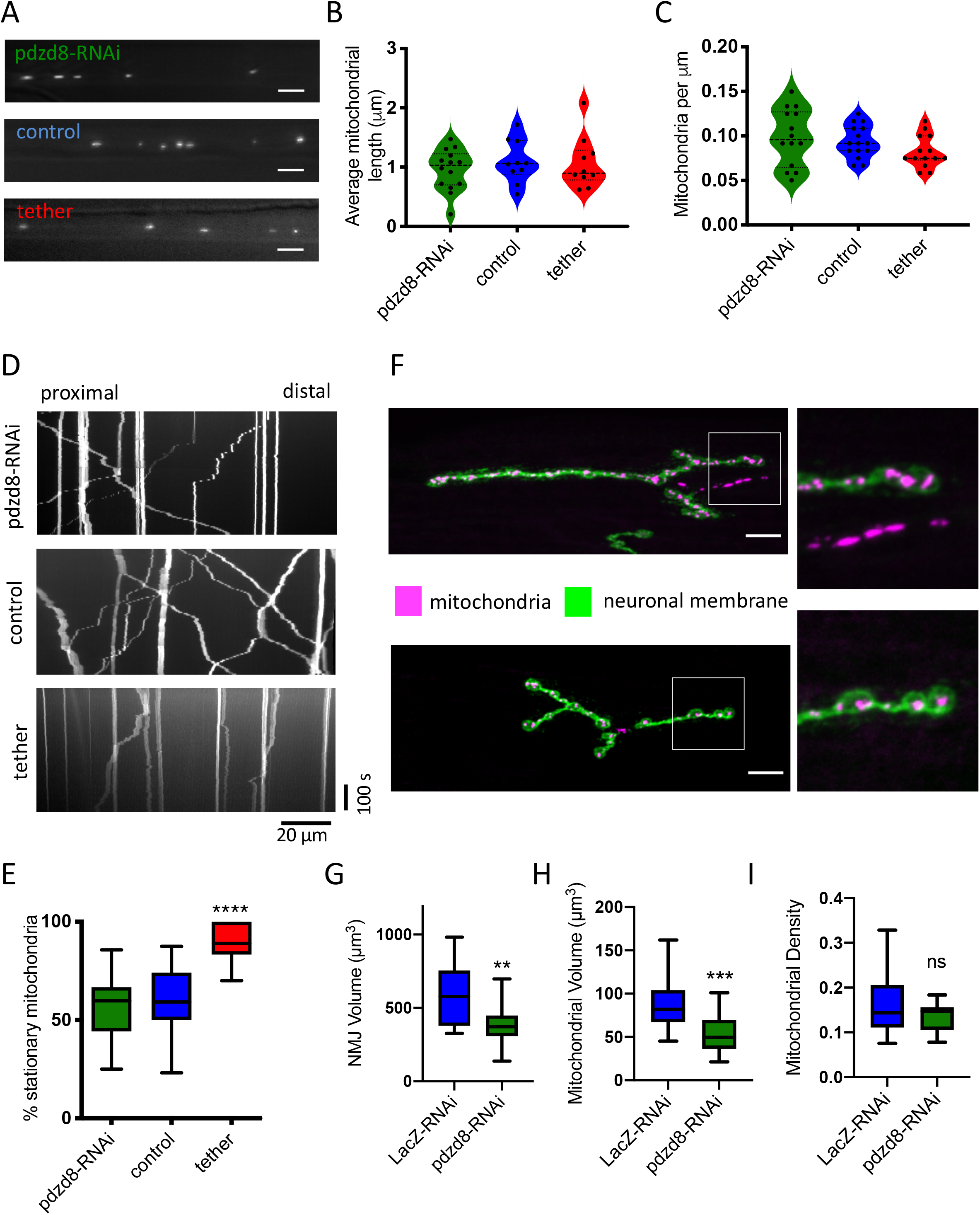
Knockdown of *pdzd8* in larval neurons causes minor defects while increasing MERCs is detrimental to in axonal mitochondria size and motility. (**A**) Representative images of mitochondrial morphology and distribution in larval axons. Mitochondria were detected using CCAP>mitoGFP in controls and *pdzd8*-RNAi expressing larvae. Scale bar 5 μm. (**B,C**) Mitochondrial length (B), and mitochondrial density (C) in the larval axons shown in A were analysed using ordinary one-way ANOVA and Holm-Sidak’s multiple comparisons. n = 10, 10, 13 animals, data points represent different axons, all differences ns. (**D**) Representative kymographs showing motility of CCAP>mitoGFP signal in controls and *pdzd8*-RNAi expressing larvae. Stationary mitochondria appear as vertical lines, moving mitochondria form diagonal lines in anterograde or retrograde directions, t = 500s (**E**) Quantification of mitochondrial transport shown in D, analysed using ordinary one-way ANOVA and Holm-Sidak’s multiple comparisons, n = 14-25, p<0.0001. (**F**) Representative images of NMJs and mitochondria of controls and *pdzd8*-RNAi labelled using OK371>mitoGFP. magenta = mitoGFP, green = anti-HRP (neuronal membrane), scale bar 10 μm. (**G-I**) Quantifications of NMJ volume, p=0.0036, (**G**), mitochondrial volume, p=0.002 (**H**) and mitochondrial density (**I**) were compared using an unpaired t-test with Welch’s Correction.

To better understand the effects of altered MERCs in neurons, we analysed the morphology of mitochondria located in 1s boutons of larval neuromuscular junctions (NMJs) on muscle 4. Knockdown of *pdzd8* led to smaller NMJs and a significant reduction of mitochondrial volume, but overall no change in mitochondrial density compared to control flies (Figure 4F-G), showing that mitochondria distribute normally in these smaller NMJs. Increased tethering, however, resulted in severely deformed NMJs (Figure S3C) and made NMJs type 1s and 1b synaptic boutons indistinguishable, making it impossible to quantify mitochondrial density in 1s boutons. Together, these results show that increasing tethering has dramatic and detrimental effects early in development but reduced tethering through *pdzd8*-RNAi expression has more limited effects during these early stages of neuronal development.

### Reducing *pdzd8* expression increases mitophagy in aged neurons

We hypothesized that the reduced sensitivity to mitochondrial toxins may be due to improved mitochondrial quality control mechanisms. Thus, we analysed the levels of mitophagy, the clearance of damaged mitochondria by autophagy, in these neurons using the mitoQC mitophagy reporter (Allen *et al*, 2013; Lee *et al*, 2018b). This reporter is a pH-sensitive mCherry-GFP fusion targeted to the mitochondrial outer membrane which provides a read out of mitophagy in the form of mCherry-only signal where the acidic lysosomal environment has quenched the GFP. Mitophagy was detected in the soma of nSyb-GAL4-expressing neurons in both larval and adult brains (Figure 5A and C). There was no significant difference in mitophagy levels in larval neurons expressing *pdzd8*-RNAi or control (Figure 5A-B). Quantification of mitoQC mCherry puncta did not indicate an age-dependent increase in mitophagy in the brains of control adults, however mitophagy was significantly increased in the brains of 20-day-old *pdzd8*-RNAi animals compared to controls (Fig 5C-D).

**Figure 5.**
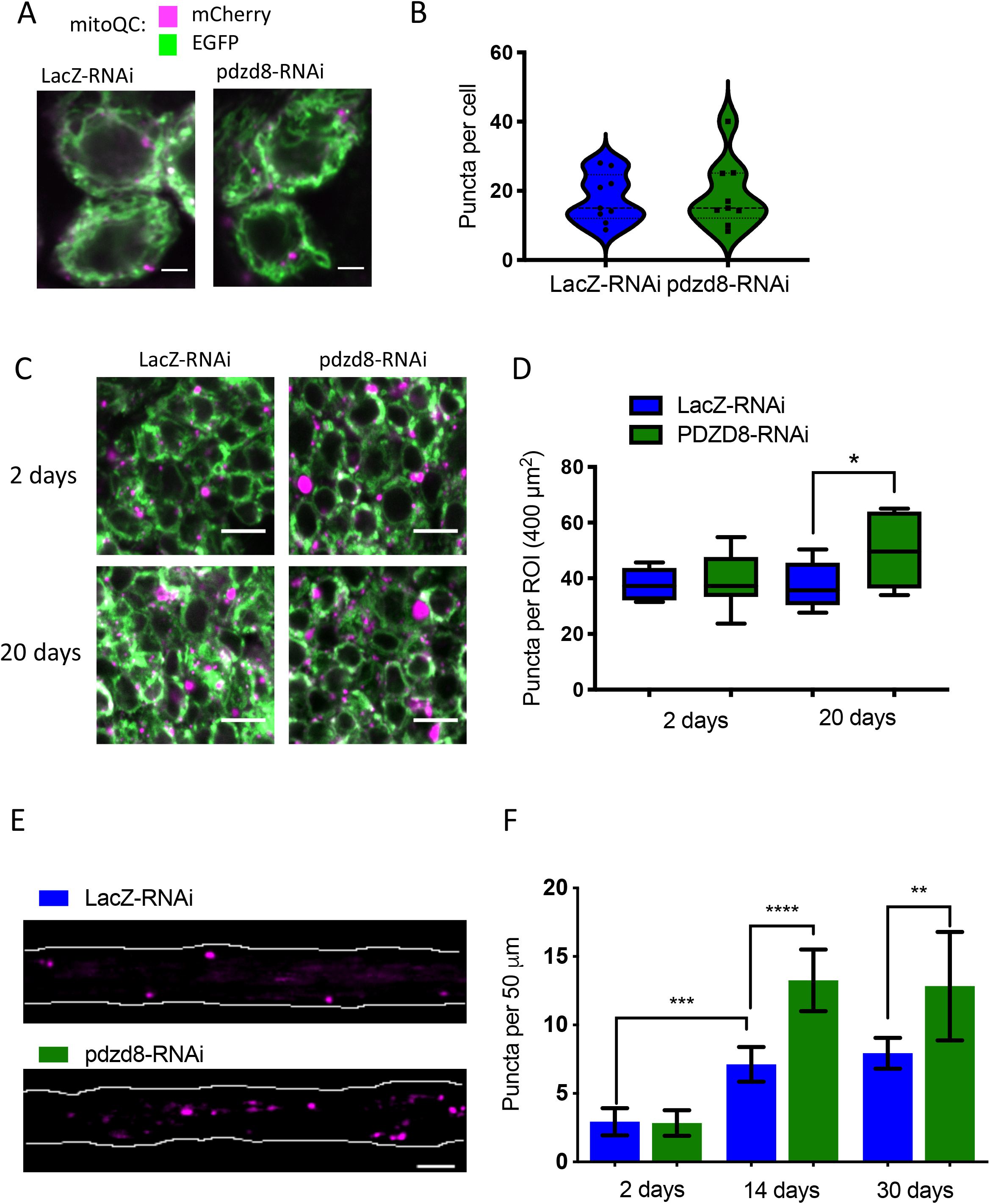
Pan-neuronal *pdzd8*-RNAi increases mitophagy during aging. (**A**) Representative images of MitoQC signal in wandering L3 larval ventral ganglia. magenta = mCherry, green = GFP, images show a single plane of a Z-stack, scale bar 2 μm. (**B**) Quantification of MitoQC puncta shown in (A), n = 9, differences ns. (**C**) Representative images of MitoQC signal in adult brains in two- and ten-day old flies, magenta = mCherry, green = GFP, scale bar 5 μm, image shows a single plane of a Z-stack. (**D**) Quantification of MitoQC signal in adult brains and compared using an unpaired t-test with Welch’s correction (n = 7-9, p=0.0072) (**E**) the Representative images of MitoQC signal in 14 day-old fly wings. Only mCherry signal (magenta) is shown for clarity, white outlines edges of wing nerves, scale bar 5 μm. (**F**) Quantification of MitoQC signal in aged fly wings at 2, 14 and 30 days post eclosion using a one-way ANOVA with Holm-Sidak’s multiple comparisons. n (2 days) = 33, 26 (14 days) = 24, 31, (30 days) = 32, 12.

The majority of mitochondria in neurons are found in the neurites, so to examine mitophagy in axons of aged flies, we analysed mitoQC signal in axons of the adult fly wing *in situ* (Vagnoni & Bullock, 2016) (Figure 5E). Here, in contrast to the adult brain cell bodies, we did observe an age dependent increase in mitophagy in axons of control flies (Figure 5F). Consistent with our previous results, *pdzd8* knockdown further increased mitophagy in axons of aged flies (Figure 5E-F). Together, these results show that loss of pdzd8 promotes the turnover of mitochondria in aging neurons.

### Reduced MERCs is protective in a fly model of Alzheimer’s disease

Our results indicate that the loss of pdzd8 is protective against mitochondrial insults, prevents the age-related decrease in locomotion and increases lifespan. Since mitochondrial dysfunction is a common feature of many neurodegenerative diseases, and altered MERCs has been documented in some, we next sought to explore the neuroprotective potential of pdzd8 depletion in an age-related neurodegenerative disease model. To test this, we turned to an Alzheimer’s disease fly model where the expression of pathogenic Aβ_42_ has been shown to cause neural dysfunction, due in part to oxidative stress (Rival *et al*, 2009).

Since increased MERCs have also been associated with Alzheimer’s disease (Area-Gomez *et al*, 2012; Del Prete *et al*, 2017), we first used the SPLICS reporter to determine the number of MERCs in larval axons. Consistent with other cellular and organismal models, flies expressing the Aβ_42_ showed an increase in SPLICS puncta, indicating MERCs are increased in axons of this model of Alzheimer’s disease (Figure 6A-B). Flies expressing Aβ_42_ in neurons exhibit a significant climbing defect in young flies that worsens rapidly with age (Figure 6C)(Crowther *et al*, 2005). However, pan-neuronal *pdzd8* knockdown in combination with Aβ_42_ was sufficient to substantially ameliorate the decline in locomotor activity observed in young and 10-day-old Aβ_42_ flies (Figure 6C). Thus, reducing pdzd8-mediated MERCs is protective in this progressive neurodegenerative disease model with increased MERCs.

**Figure 6.**
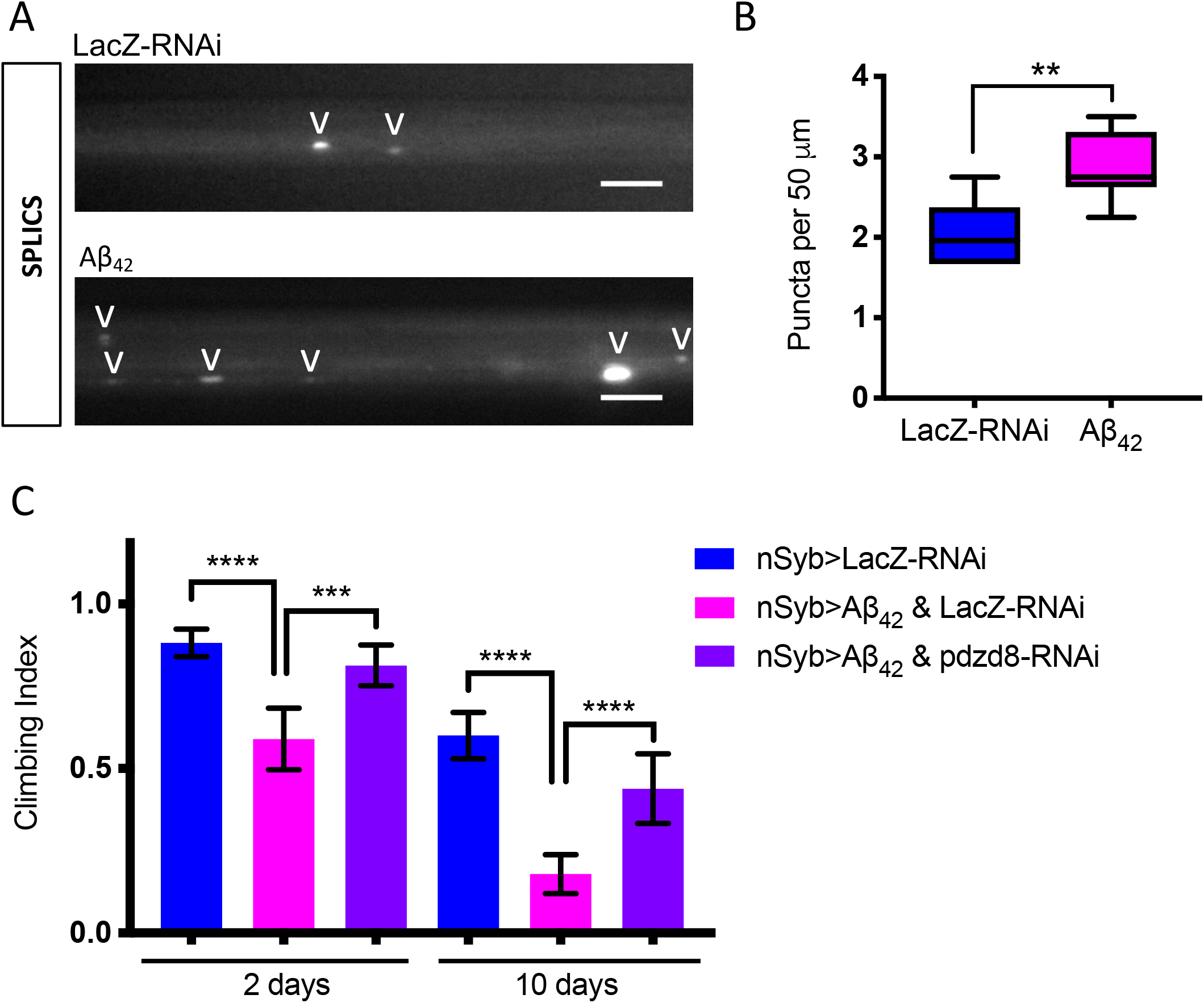
Reducing pdzd8-mediated MERCs rescues the locomotor defects in an Alzheimer’s disease *Drosophila* model. (**A**) CCAP>SPLICS signal in larval axons expressing Aβ_42_ compared to controls. Quantified puncta highlighted with V. (**B**) Quantification of SPLICS signal in (A) using an unpaired t-test with Welch’s correction, n = 6, p=0.008. (**C**) Aged climbing assay of flies expressing nSyb>Aβ_42_, n>75 flies were aged and climbed on the indicated days. p <0.001, 0.002, <0.001, <0.001.

## Discussion

Here we have identified and characterized the putative *Drosophila* homolog of the newly discovered mammalian MERC tethering protein Pdzd8 (Hirabayashi *et al*, 2017). The sequence divergence between Pdzd8 and its yeast paralog Mmm1 and the additional domains present in Pdzd8 (Wong & Levine, 2017), made the relationship between these paralogs difficult to identify (Wideman *et al*, 2018). While the conserved predicted domain structure strongly suggests *Drosophila CG10362* encodes the fly homolog of mouse Pdzd8, the low, 21 % overall sequence identity of these proteins raises the interesting possibility of evolution of more species-specific functional specialisation.

Using RNAi, we characterized the effects of depletion of pdzd8 in *Drosophila* with a focus on neurons where this protein is most highly expressed. Knockdown of *pdzd8* reduces contacts between the ER and mitochondria in epidermal cells measured using super resolution microscopy of ER and mitochondria labeled with fluorescent reporters, in motor neurons, monitored by the contact site reporter SPLICS, and in the soma of adult neurons using transmission electron microscopy. These data suggest that pdzd8, like its mammalian ortholog, functions as a tether between ER and mitochondria. The only other neuronally expressed tethering protein that has been characterized in flies is the *Drosophila* ortholog of Mfn2, Marf (Hwa *et al*, 2002). However, the analysis of Marf is complicated by its additional roles in mitochondrial and ER morphology (Debattisti *et al*, 2014; El Fissi *et al*, 2018; Sandoval *et al*, 2014). While mammalian Pdzd8 is expressed in a range of tissues (Hirabayashi *et al*, 2017), in flies,*pdzd8* mRNA expression is low outside the nervous system. Knockdown of *pdzd8* and expression of a synthetic tether therefore provided a unique opportunity to simply and selectively examine the function of MERCs in *Drosophila* neurons.

We found increased ER-mitochondrial tethering in neurons strongly impairs climbing ability and reduces lifespan of flies. Such detrimental effects are consistent with previous reports using this line (Basso *et al*, 2018). Other similar manipulations have also been shown to result in dopaminergic neuron loss (Lee *et al*, 2018c) and detrimental effects on sleep in ventral lateral neurons (Valadas *et al*, 2018). Here we also show highly abnormal NMJ development, with increased MERCs and smaller, but otherwise structurally intact NMJs upon *pdzd8* knockdown. The function of the yeast paralog Mmm1 and of its Synaptotagmin-like mitochondrial lipid-binding protein (SMP) domains, suggests that disruptions in lipid transfer due to less pdzd8 might contribute to these developmental defects (Jeong *et al*, 2017; Kawano *et al*, 2018; Shirane *et al*, 2020), however whether lipid biogenesis defects could contribute to a protective effect of *pdzd8*-RNAi in neurons remains an open question.

Transport of mitochondria is intimately linked to the health of neurons, at least in the peripheral nervous system (De Vos *et al*, 2008; Harbauer, 2017; Misgeld & Schwarz, 2017). Although there is some evidence that increased MERCs may be directly associated with decreased mitochondrial motility (Krols *et al*, 2018), this has not been shown in neurons which are particularly sensitive to mitochondrial transport imbalance (Maday *et al*, 2014). Our data suggests axonal mitochondrial transport defect contributes to the detrimental effects of increased tethering and adds to the evidence that efficient mitochondrial transport is essential for healthy aging neurons, as seen in many models of neurodegenerative motor disorders (Baldwin *et al*, 2016).

In contrast to the detrimental effects of increased tethering, reducing MERCs by knockdown of *pdzd8* in *Drosophila* neurons dramatically delayed age-associated decline in locomotor activity and significantly extended median lifespan compared to control animals. Since decline in mitochondrial transport is proposed to contribute to neuronal aging (Vagnoni & Bullock, 2018), we hypothesized that reducing tethering in the aging flies might be protective by allowing sustained mitochondrial motility (Mattedi & Vagnoni, 2019). However, we detected no change in the percentage of motile mitochondria in larval neurons with reduced *pdzd8* expression. Since knockdown of *pdzd8* also prolonged the survival of flies fed mitochondrial toxins, this suggests that the protective effects of reducing MERCs might instead be result from more efficient clearance of the damaged mitochondria accumulating with age or more acutely by feeding flies mitochondrial toxins.

To explore the clearance of damaged mitochondria, we examined the age dependence of mitophagy using the mitoQC reporter. Clearance of damaged cellular components by autophagy is critical for the maintenance of healthy neurons with age (Stavoe & Holzbaur, 2019). More specifically, the clearance of damaged mitochondria via mitophagy is also thought to be a key factor in healthy aging of neurons (Ma *et al*, 2018; Pickrell & Youle, 2015; Whitworth & Pallanck, 2017). It remains unclear, however whether, increased or decreased mitophagy in neurons is protective during aging (Montava-Garriga & Ganley, 2020) and if specific neuronal populations are more susceptible to mitochondrial turnover. Studies examining mitophagy in aging brains have produced inconsistent results, with differences observed between brain regions and cell types (Cornelissen *et al*, 2018; Lee *et al*, 2018b; Sun *et al*, 2015). Consistent with our previous result in dopaminergic neurons (Lee *et al*, 2018b), we also did not detect any age-related change in mitophagy levels in the cell bodies in the brains of control flies.

There is accumulating evidence showing that axonal mitophagy occurs in cultured mammalian neurons (Ashrafi *et al*, 2014; Zheng *et al*, 2019), but other results suggest axonal maintenance in flies may not require mitophagy (Cao *et al*, 2017). There is evidence of mitophagy *in vivo* in neurites in young mice (McWilliams *et al*, 2018; McWilliams *et al*, 2019), but mitoQC has not previously been used to detect mitophagy in neurites during aging (Cornelissen *et al*, 2018). Using mitoQC, we detected a significant amount of mitophagy in wing axons of control flies which increased with age. Our results suggest mitophagy in axon but not cell bodies increases with age, but future experiments should specifically examine variations in mitophagy with age across brain regions, cell types and even other subcellular compartments such as dendrites.

Here we provide the first *in vivo* evidence that mitophagy may be regulated by pdzd8-mediated MERCs. Mitophagy levels did not change in young flies with less pdzd8, but when aged, these flies displayed significant increases in mitophagy in both soma and axons. There is also accumulating evidence that defects in mitophagy contribute to the early stages of Alzheimer’s disease (AD) (Kerr *et al*, 2017; Lee *et al*, 2019) and boosting mitophagy has been shown to be protective in worm and mouse AD models (Fang *et al*, 2019). Here we show increased MERCs in an AD fly model consistent with the altered MERCs identified in both patients and cell models of AD (Area-Gomez *et al*, 2012). Expression of *pdzd8*-RNAi in the AD flies significantly slowed their age-associated decline in climbing, consistent with reduced MERCs allowing more efficient mitophagy and removal of damaged mitochondria.

Mitochondrial dysfunction and associated metabolic changes are considered a hallmark of aging (Lopez-Otin *et al*, 2013). Boosting mitophagy has been suggested to increase lifespan and protect animals from mitochondrial toxins in both *D. melanogaster* (Aparicio *et al*, 2019; Rana *et al*, 2017) and *C. elegans* (Palikaras *et al*, 2015). Early work on starvation induced autophagosome biogenesis showed MERCs could contribute to the formation of both the autophagosome (Hailey *et al*, 2010; Hamasaki *et al*, 2013) and the mitophagosome (Yang & Yang, 2013). As postmitotic, long-lived cells with complex morphologies, neurons require more careful regulation of mitophagy than other cell types (Evans & Holzbaur, 2019), but the mechanisms that regulate this process in neurons are still poorly understood (Evans & Holzbaur, 2020).

MERCs mediated by pdzd8 may limit the rate of mitophagy, analogous to the protective role that mitochondrial fusion is thought to play during starvation-induced autophagy (Gomes *et al*, 2011; Rambold *et al*, 2011; Rana *et al*, 2017; Twig *et al*, 2008). Recently it was reported that increasing several different types of MERC proteins can slow toxin induced mitophagy in non-neuronal cultured cells (McLelland & Fon, 2018; McLelland *et al*, 2018). There is also some recent evidence that MERCs might be directly involved in regulation of mitophagy in neurons (Puri *et al*, 2019). Pdzd8 has also been found to mediate contacts between the ER and lysosomes (Guillen-Samander *et al*, 2019) and three way contacts between the ER, mitochondria and late endosomes (Elbaz-Alon *et al*, 2020; Shirane *et al*, 2020). This may indicate that Pdzd8, like other MERC proteins, can play multiple roles at different organelle contacts, or that Pdzd8 is functional at three-way contacts between the ER, lysosomes and mitochondria (Wong *et al*, 2019). Pdzd8 function at any of these contact sites could potentially alter mitophagy and future work should explore the mechanisms behind our observed increases in mitophagy.

While our data suggests that increased mitophagy contributes to the protective effects of the loss of pdzd8-mediated MERCs, we have not ruled out contributions from the other specialized functional and signalling roles of MERCs in metabolic regulation, lipid biogenesis and calcium handling (Lee *et al*, 2018a; Rowland & Voeltz, 2012). The increased mitophagy we detected in aged neurons expressing *pdzd8*-RNAi correlates with improved survival and aged locomotor activity, suggesting that reducing the extent of pdzd8-mediated MERCs may facilitate more efficient mitophagy and removal of damaged mitochondria from aging neurons. We propose that reducing pdzd8-mediated MERCs may be protective in aging neurons by allowing more efficient turnover of mitochondria as damage accumulates with age. As regulators of mitophagy, manipulating MERCs may provide an avenue for enhanced mitochondrial quality control to help promote healthy aging of neurons.

## Methods

### Husbandry

Flies were raised under standard conditions at 25°C on food containing agar, cornmeal, molasses, malt extract, soya powder, propionic acid, nipagin and yeast in a 12 h:12 h light:dark cycle.

### Genetics

*Drosophila* lines were obtained as indicated in Table 1, or generated as described below. All mutant lines used in this study were backcrossed to an isogenic *w*^1118^ strain (RRID:BDSC_6326), for 4-6 generations before use. For all integration events, multiple independent lines were initially isolated, verified by PCR and assessed for consistent effects before selecting a single line of each integration site for further study. Wherever possible inert UAS lines such as UAS-*LacZ*-RNAi and UAS-mitoCherry are used as dilution controls to ensure equal numbers of UAS constructs in control and experimental conditions. Unless otherwise stated male flies were used in all experiments.

### New lines

#### SPLICS

The SPLICSs construct with an 8-10 nm range from Tito Cali & Marisa Brini (Cieri *et al*, 2017), was amplified from pSYC-SPLICSs-P2A using (TAAGCAGCGGCCGCTGATTTAGGTGACACTATAG) and T7 forward primer (TAATACGACTCACTATAGGG) and cloned into pUAST-AttB between NotI and XbaI sites. Flies were injected by BestGene to insert into attP16 (II) and attP2 (8622, III) and the attP16 site gave better signal and so was used in this work. The number of puncta produced in the axon bundles driven by nSyb-GAL4 varied with a consistently more puncta than in the central axons bundles in the peripheral bundles (Fig S2B).

#### pdzd8-HA

pdzd8-HA was synthesized by Genewiz based on the cDNA Genbank sequence LD34222 (AY118553.1) (Sayers *et al*, 2020), and cloned into pUAST.attB between EcoRI and XbaI. The University of Cambridge Department of Genetics Fly Facility generated lines by injection of this construct into the attP40 landing site.

### Climbing

The startle induced negative geotaxis (climbing/locomotor) assay was performed as described previously (Andreazza *et al*, 2019). Briefly, a maximum of 23 males were placed into the first tube of a countercurrent apparatus, tapped to the bottom, and given 10 s to climb 10 cm. This procedure was repeated five times (five tubes), and the number of flies that has remained into each tube counted and the climbing performance expressed as a climbing index (Greene *et al*, 2003). The same flies were aged and assayed again on the indicated days post-eclosion.

### Lifespan

For lifespan experiments groups of approximately 20-25 males were collected with minimal time under anesthesia (CO_2_), placed into separate vials with food and maintained at 25°C. Flies were transferred into fresh vials every 2-3 days, and the number of dead flies were recorded. Percent survival was calculated using https://flies.shinyapps.io/Rflies/. To assess lifespan in a diet with restricted nutrients flies were raised in standard conditions then transferred to tubes containing food made from 5 % sucrose and 1 % agar and flipped every 2-3 days. Lifespans in the presence of mitochondrial toxins and hydrogen peroxide were also performed on food made from 5 % sucrose and 1 % agar cooled to less than 50°C before adding toxin at 1:1000. Rotenone (Sigma R8875) was dissolved in DMSO (1 mM final concentration) and antimycin A (Sigma A8674) (4 μg/mL final concentration) dissolved in 70 % ethanol. Flies in toxin assays were starved for 5 h before being placed on food containing toxins. Flies in rotenone assays were monitored twice a day and flipped every two days. Flies in antimycin A assays were monitored three times a day and flipped every two days.

### Fluorescence Microscopy

Imaging of larval axons was performed as described in (Wang & Schwarz, 2009) with the following variations: wandering third instar larvae were pinned at each end dorsal side up to a reusable Sylgard (Sigma 761028) coated slide using pins (Fine Science Tools FST26002-10) cut to ~5 mm and bent at 90°. The larvae were cut along the dorsal midline using micro-dissection scissors. Internal organs were removed with forceps without disturbing the ventral ganglion and motor neurons. Larvae were then covered in dissection solution (Godena *et al*, 2014). The cuticle was then pulled back with four additional pins. The anterior pin was adjusted to ensure axons are taut and as flat as possible for optimal image quality.

Movies were taken using a Nikon E800 microscope with a 60x water immersion lens (NA 1.0 Nikon Fluor WD 2.0) and an LED light source driven by Micromanager 1.4.22 Freeware (Edelstein *et al*, 2014). A CMOS camera (01-OPTIMOS-F-M-16-C) was used to record 100 frames at a rate of 1 frame per 5 s (8 min 20 s total). Axons were imaged within 200 μm of the ventral ganglion in the proximal portion of the axons and no longer than 1 h after dissection. Movies were converted into kymographs using Fiji (Schindelin *et al*, 2012) and mitochondrial motility quantified manually with the experimenter blinded to the condition.

For SPLICS imaging in axon bundles at least three ROI 50 μm x 12 μm were quantified per animal and averages for each larva were plotted. For SPLICS quantification puncta intensity varied considerably so blinded manual counting was used.

To image NMJs larvae were dissected as described above and fixed for 20 min in 4 % formaldehyde in PBS. After blocking for 1 h in 1 % BSA/0.3 % Triton X-100/PBS solution, anti-HRP was added at 1:500 and samples agitated gently overnight at 4°C. After three washes in 0.3 % Triton X-100/PBS at room temperature, samples were incubated with Alexa Fluor 594 at 1:500 for 1 h in 1% BSA/0.3% Triton X-100/PBS solution. Samples were then washed in 3x in PBS before being mounted in Prolong Diamond. NMJs were imaged on a Nikon Eclipse TiE inverted microscope with appropriate lasers using an Andor Dragonfly 500 confocal spinning disk system, using an iXon Ultra 888 EMCCD camera (Andor), coupled with Fusion software (Andor) using a 60x NA 1.49 objective. NMJs on muscle 4 from segments A3 and A4 (NMJs on these segments are the same size (Nijhof *et al*, 2016)) were captured in Z stacks with 0.3 μm step size and analysed using Imaris (x64 9.2.0) to determine NMJ volume, mitochondrial volume and mitochondrial number.

For MitoQC imaging, samples were fixed for 30 min in 4 % formaldehyde (16 % VWR 100503) diluted in pH 7.0 PBS. Adult brains were mounted in Prolong Diamond Antifade Mountant (Thermofisher P36961) using spacers and imaged on a Carl Zeiss LSM880 confocal laser-scanning system on an Axio Observer Z1 microscope (Carl Zeiss), coupled with ZEN software (Carl Zeiss) using a 100x Plan-APOCHROMAT /1.4 oil DIC objective. Images are shown in false colour with magenta puncta representing mCherry signal indicating where the reporter is in an acidic environment of a lysosome and the GFP has been quenched (Allen *et al*, 2013).

Imaging of wings was performed as described in Vagnoni & Bullock (Vagnoni & Bullock, 2016). Briefly, flies were anaesthetized with CO_2_, and immobilised with their wings outstretched on a cover glass with a fine layer of Halocarbon oil (VWR). A second cover glass was then added on top of the fly to stabilize the sample. Live imaging in the wing nerves was performed using a Nikon spinning disk system essentially as described previously (Morotz *et al*, 2019). The mitoQC puncta were annotated with Fiji using the Cell Counter plugin and quantified with the experimenter blinded to the genotype.

For imaging of the larval epidermal cells, the larvae were dissected as described above but the nervous system was also removed before fixation. The samples were washed in PBS and the muscles were then removed (Tenenbaum & Gavis, 2016). The dissected filets were mounted in Prolong Diamond Antifade Mountant using No. 1.5H High Precision Deckglaser cover slips and placed under a weight for 24 h. N-SIM (Nikon Structured Illumination Microscopy) imaging was performed on a Nikon Ti Eclipse with an Andor DU-897 X-5835 camera and SR Apo TIRF 100x (NA1.5) objective run using NIS-Elements 4.60. Images were analysed in Fiji (Schindelin *et al*, 2012) using the Coloc2 plugin.

### Transmission Electron Microscopy

Transmission electron microscopy (TEM) was performed at Cambridge Advanced Imaging Center (CAIC). Brains of 2 day old adult flies were fixed in 2 % glutaraldehyde/ 2 % formaldehyde in 0.1 M sodium cacodylate buffer pH 7.4 containing 2 mM CaCl_2_ and 0.1 % Tween20 (based on method described in (Celardo *et al*, 2016)), overnight at 4°C. Samples were then washed 5x with 0.1 M sodium cacodylate buffer and then treated with osmium for 2 days at 4°C (1 % OsO_4_, 1.5 % potassium ferricyanide, 0.1 M sodium cacodylate buffer pH 7.4). Samples were then washed 5x in distilled water and treated with 0.1 % aqueous thiocarbohydrazide for 20 min in the dark at room temperature. Samples were washed another 5x in distilled water then treated with osmium a second time for 1 h at room (2 % OsO_4_ in distilled water). Samples were then washed another 5x in distilled water before being treated with uranyl acetate bulk stain for 3 days at 4°C (2 % uranyl acetate in 0.05 M maleate buffer pH 5.5). After a final 5x wash in distilled water, samples were dehydrated in 50/70/95/100 % ethanol, 3x in each for at least 5 min each. Dehydration was completed by two further treatments with 100 % dry ethanol, 2x in 100 % dry acetone and 3x in dry acetonitrile for at least 5 min each. Quetol resin mix (12 g Quetol 651, 15.7 g NSA, 5.7 g MNA, 0.5 g benzyldimethylamine) made with an equal volume of 100 % dry acetonitrile and samples placed in this mix for 2 h at room temperature. Samples were then incubated in pure Quetol resin mix for 5 days, exchanging the samples to fresh resin mix each day. After 5 days the brains were embedded in coffin moulds and cured at 60°C for at least 48 h. Ultrathin sections were cut on a Leica Ultracut E at 70 nm thickness. Sections were mounted on 400 mesh bare copper grids and viewed in a FEI Tecnai G20 electron microscope run at 200 keV using a 20 μm objective aperture. Quantification of the percentage of clearly identifiable mitochondria in contact with ER was performed manually as described in (Celardo *et al*, 2016) and the experimenter was blinded to the genotype.

### qPCR

Five female wandering third instar larvae per sample were washed briefly in 1xPBS, placed in RNAse free tubes and frozen on dry ice. Larvae were homogenized in Trizol and RNA isolated by phenol:chloroform extraction and isopropanol precipitation. DNAse treatment using Invitrogen TURBO DNA-free rigorous procedure was performed before measuring RNA concentration with a Qubit^®^ RNA HS Assay Kit (Molecular Probes, Life Technologies). Reverse transcription reactions used 1.32 μg of RNA using SuperScript III Reverse Transcriptase (Invitrogen) with Oligo(dT)23VN as per manufacturer’s instructions. The resulting cDNA was used for qPCRs using PowerUp SybrGreen (Applied Biosystems A25742). Primers for pdzd8 were PDZD8-F TTCTGTTTGGCTTCTCCTGG, PDZD8-R TTGAGGAACTGCGACTGATC designed using RealTime qPCR Assay Entry (idtdna.com). *αTub84B* (Fwd: TGGGCCCGTCTGGACCACAA, Rev: TCGCCGTCACCGGAGTCCAT), *vkg* (Fwd: CGAGGATGTTACCCAGAGATC, Rev: TGCGTCCCTTGATTCCTTTG), *COX8* (Fwd: CAGAGCCGTTGCCAGTC, Rev: CTTGTCGCCCTTGTAGTCC), and *Rpl32* (Fwd: AAACGCGGTTCTGCATGAG, Rev: GCCGCTTCAAGGGACAGTATCTG) were used as housekeeping genes with their values combined to compare knockdown to the geometric mean (Taylor *et al*, 2019).

### ATP

ATP levels were measured in two- and twenty-day-old fly heads with 40 flies per genotype and three biological replicates. The ATP levels were measured as described in Tufi, Gleeson *et al*. (Tufi *et al*, 2019) with minor modifications. Briefly, heads were homogenized in 6 M guanidine-Tris/EDTA extraction buffer and subjected to rapid freezing in liquid nitrogen. Luminescence produced from homogenates mixed with the CellTiter-Glo Luminescent Cell Viability Assay (Promega) was measured with a SpectraMax Gemini XPS luminometer (Molecular Devices) and normalized to total protein, quantified using the Pierce BCA method (Thermo Fisher Scientific).

### Quantification & Statistical analysis

Statistical analyses were performed using GraphPad Prism 8 software. Data are reported as mean ± 95 % CI unless otherwise stated in figure legends. Climbing was assessed using a Kruskal-Wallis non-parametric test with Dunn’s post-hoc correction for multiple comparisons. Lifespans were compared using log-rank Mantel-Cox tests. Number of flies and p-values are reported in the figure legends.

Mitochondrial transport was analysed using ordinary one-way ANOVA and Holm-Sidak’s multiple comparison. ATP measurements were analysed by two-tailed t-test. Values are not significantly different to controls unless otherwise stated.

SCope (http://scope.aertslab.org/) was used to visualize transcriptome data from the unfiltered adult fly brain dataset (Davie *et al*, 2018).

---

**Table S1:**
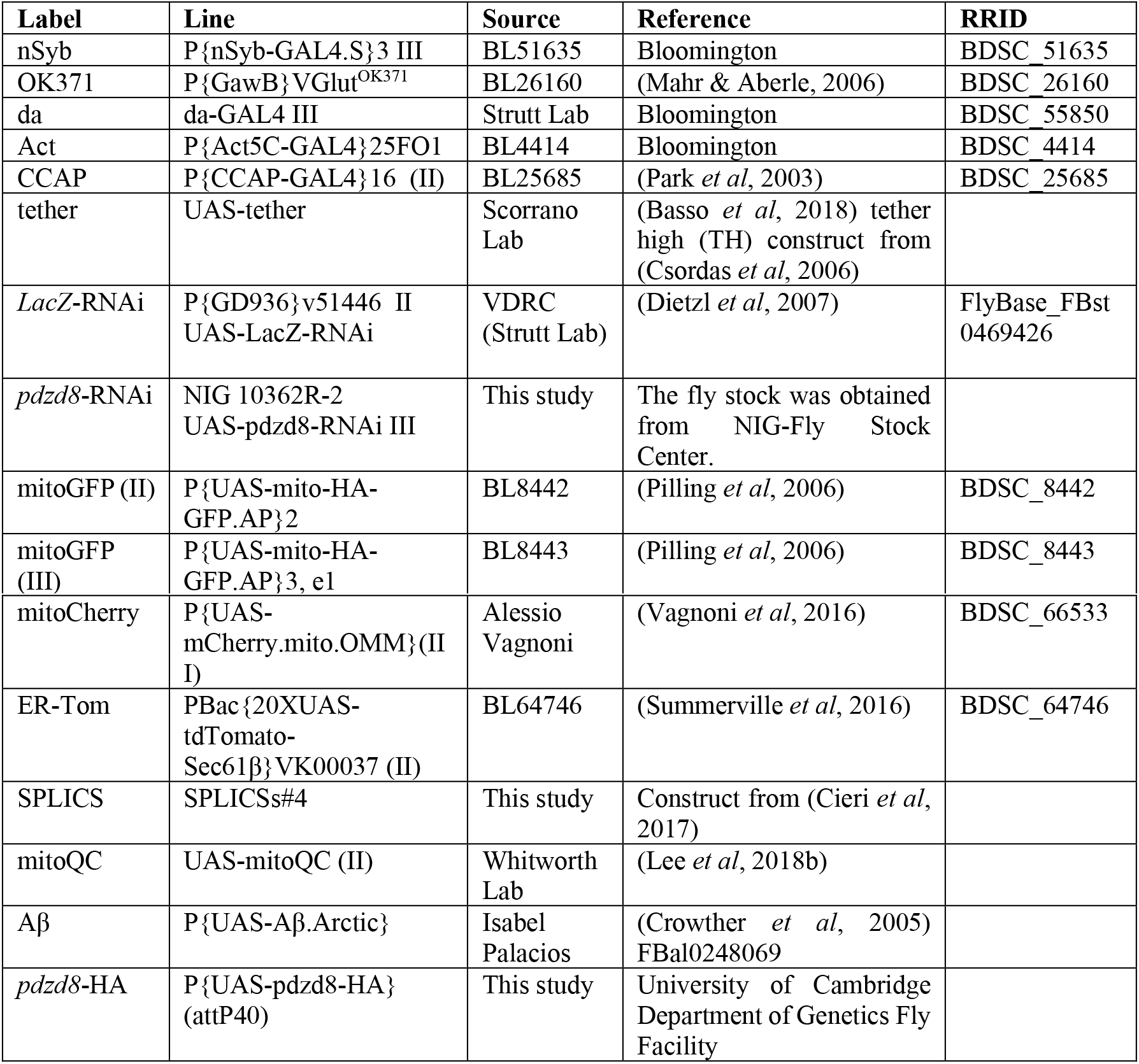
Fly Lines

| Label | Line | Source | Reference | RRID |
| --- | --- | --- | --- | --- |
| nSyb | P{nSyb-GAL4.S}3 III | BL51635 | Bloomington | BDSC_51635 |
| OK371 | P{GawB}VGlut<sup>OK371</sup> | BL26160 | (Mahr & Aberle, 2006) | BDSC_26160 |
| da | da-GAL4 III | Strutt Lab | Bloomington | BDSC_55850 |
| Act | P{Act5C-GAL4}25FO1 | BL4414 | Bloomington | BDSC_4414 |
| CCAP | P{CCAP-GAL4}16 (II) | BL25685 | (Park *et al*, 2003) | BDSC_25685 |
| tether | UAS-tether | Scorrano Lab | (Basso *et al*, 2018) tether high (TH) construct from (Csordas *et al*, 2006) |  |
| LacZ-RNAi | P{GD936}v51446 II<br>UAS-LacZ-RNAi | VDRC (Strutt Lab) | (Dietzl *et al*, 2007) | FlyBase_FBst_0469426 |
| pdzd8-RNAi | NIG 10362R-2<br>UAS-pdzd8-RNAi III | This study | The fly stock was obtained from NIG-Fly Stock Center. |  |
| mitoGFP (II) | P{UAS-mito-HA-GFP.AP}2 | BL8442 | (Pilling *et al*, 2006) | BDSC_8442 |
| mitoGFP (III) | P{UAS-mito-HA-GFP.AP}3, e1 | BL8443 | (Pilling *et al*, 2006) | BDSC_8443 |
| mitoCherry | P{UAS-mCherry.mito.OMM}(II I) | Alessio Vagnoni | (Vagnoni *et al*, 2016) | BDSC_66533 |
| ER-Tom | PBac{20XUAS-tdTomato-Sec61β}VK00037 (II) | BL64746 | (Summerville *et al*, 2016) | BDSC_64746 |
| SPLICS | SPLICSs#4 | This study | Construct from (Cieri *et al*, 2017) |  |
| mitoQC | UAS-mitoQC (II) | Whitworth Lab | (Lee *et al*, 2018b) |  |
| Aβ | P{UAS-Aβ.Arctic} | Isabel Palacios | (Crowther *et al*, 2005)<br>FBal0248069 |  |
| *pdzd8*-HA | P{UAS-*pdzd8*-HA}(attP40) | This study | University of Cambridge<br>Department of Genetics Fly Facility |  |

**Table S2:** Genotypes in figures

|  |  |
| --- | --- |
| Figure 1 (B, C) | *LacZ*-RNAi/+; nSyb/+<br>*pdzd8*-RNAi/nSyb |
| Figure 1 (D, E) | SPLICS/*LacZ*-RNAi; nSyb/+<br>SPLICS/+; *pdzd8*-RNAi/nSyb |
| Figure 1 (F, G) | ER-Tom, mitoGFP/ *LacZ*-RNAi; da/+<br>ER-tom, mitoGFP/+; da/*pdzd8*-RNAi |
| Figure 2 (A, C) | *LacZ*-RNAi/+; nSyb/+<br>*pdzd8*-RNAi/nSyb |
| Figure 2 (B, D) | *LacZ*-RNAi/+; nSyb/+<br>tether/nSyb |
| Figure 3 (A-D) | *LacZ*-RNAi/+; nSyb/+<br>*pdzd8*-RNAi/nSyb |
| Figure 4 (A-E) | CCAP/+; mitoGFP/+<br>CCAP/+; *pdzd8*-RNAi/mitoGFP<br>CCAP/+; tether/mitoGFP |
| Figure 4 (F-I) | OK371, mitoGFP/*LacZ*-RNAi<br>OK371, mitoGFP/+; *pdzd8*-RNAi/+ |
| Figure 5 (A-F) | mitoQC/*LacZ*-RNAi; nSyb/+<br>mitoQC/+; nSyb/*pdzd8*-RNAi |
| Figure 6 (A-B) | *LacZ*-RNAi/SPLICS; nSyb/+<br>Aβ/SPLICS; nSyb/+ |
| Figure 6 (C) | *LacZ*-RNAi/+; nSyb/+<br>*LacZ*-RNAi/Aβ; nSyb/+<br>*LacZ*-RNAi/Aβ; *pdzd8*-RNAi/nSyb |
| Figure S1 (C) | Act/*LacZ*-RNAi (females)<br>Act/+; *pdzd8*-RNAi/+ (females) |
| Figure S1 (D) | ER-Tom, mitoGFP/*LacZ*-RNAi, nSyb/+<br>ER-Tom, mitoGFP/+; nSyb/*pdzd8*-RNAi |
| Figure S1 (E) | ER-Tom, mitoGFP/ *LacZ*-RNAi; da/+<br>ER-tom, mitoGFP/+; da/ *pdzd8*-RNAi |
| Figure S2 (B) | SPLICSs/+; nSyb-GAL4/+ |
| Figure S2 (D) | SPLICS/*LacZ*-RNAi; nSyb/+<br>SPLICS/+; tether/nSyb |
| Figure S3 (A) | OK371/*LacZ*-RNAi |
|  | OK371/+; *pdzd8*-RNAi/+ |
| Figure S3 (B) | *LacZ*-RNAi/+; mitoCherry/nSyb, |
|  | *LacZ*-RNAi/*pdzd8*-HA; nSyb/+ |
|  | *pdzd8*-HA/+; *pdzd8*-RNAi/nSyb |
|  | tether/nSyb |

**Figure S1.**
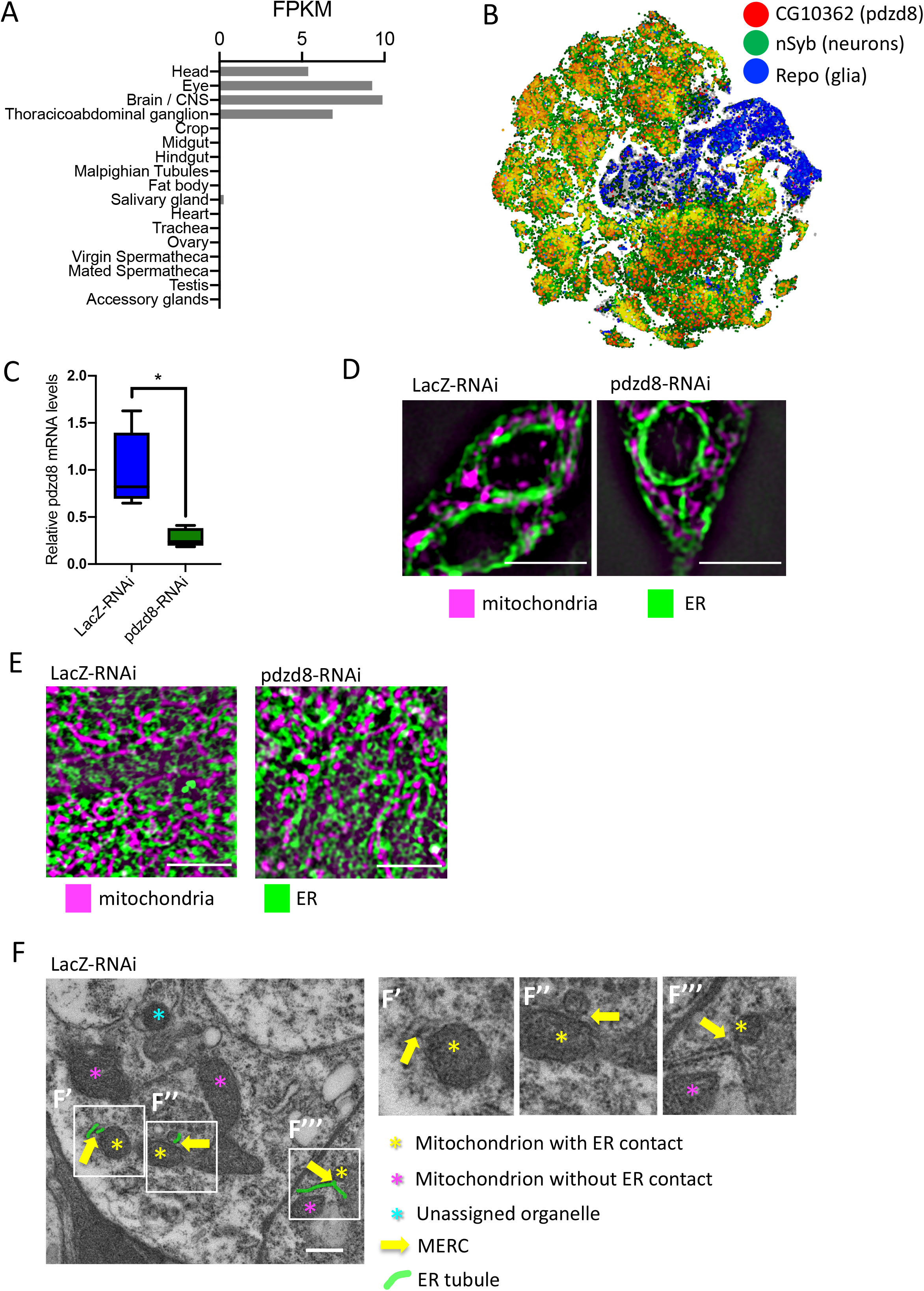
Expression and tissue specificity of pdzd8. (**A**) Tissue specific expression of pdzd8 in 7 day old adult males from (Leader *et al*, 2018) FPKM: fragments per kilobase of exon model per million reads mapped providing a normalized estimation of gene expression based on RNA-seq data. (**B**) SCope transcriptome data from the unfiltered adult fly brain dataset. (**C**) Relative abundance of *pdzd8* transcript in controls compared to Tub>*pdzd8*-RNAi normalised to the relative to geometric mean of housekeeping genes *αTub84B, vkg, COX8* and *Rpl32;* p=0.0134. (**D**) Representative SIM images of ER (green) and mitochondria (purple) in L3 larval neurons, scale bar 5 μm. (**E**) Representative SIM images of ER and mitochondria in larval epidermal cells, scale bar 5 μm. (**F**) Close up inlays (F’-F’’’) of example MERCs in Figure 1E identified in nSyb>*LacZ*-RNAi brains imaged using electron microscopy. Scale bar 500 nm.

**Figure S2.**
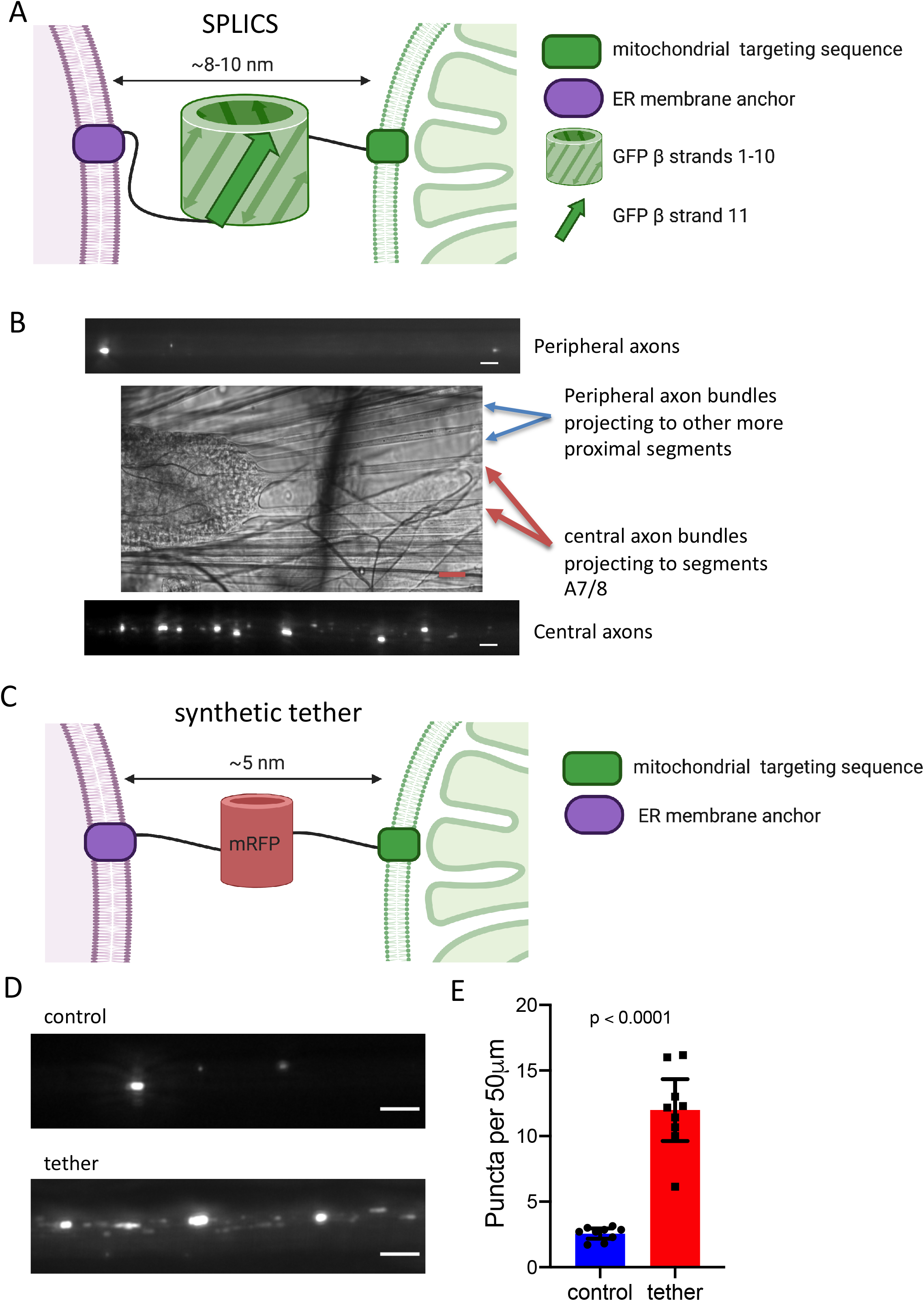
SPLICS and tether constructs used in this study. (**A**) Cartoon of SPLICS targeting and mode of action (Created with BioRender.com). (**B**) Density of SPLICS puncta in axons is different in different axon bundles, scale bar 5 μm. (**C**) Cartoon of the synthetic tether construct targeting and mode of action (Created with BioRender.com). (**D**) nSyb>SPLICS puncta indicating contact sites in larval axons, scale bar 5 μm, n = 9 per genotype.

**Figure S3.**
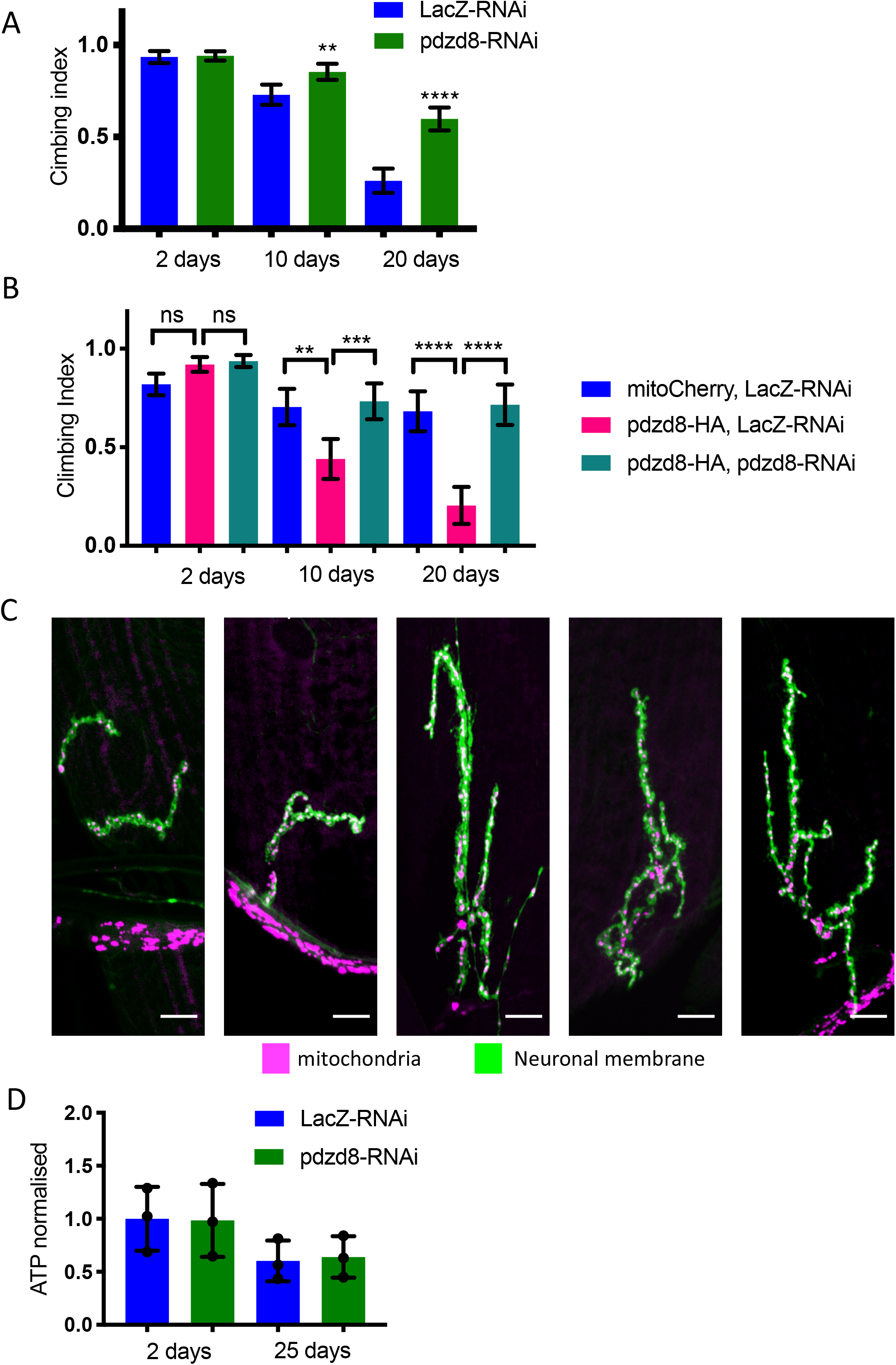
Phenotypic characterization of altered tethering in motor neurons. (**A**) Motor neuron-specific aged climbing assay showing OK371>*pdzd8*-RNAi compared to *LacZ-* RNAi controls. (**B**) *pdzd8*-RNAi rescues aged climbing defect resulting from *pdzd8* over expression. (**C**) nSyb>tether expression resulted in severely deformed NMJs on muscle 4 and made it impossible to distinguish NMJs type 1s and 1b synaptic boutons (therefore mitochondrial density could not be quantified). (**D**) ATP levels in fly heads analysed showing mean ± standard deviation, n = 3, 40 flies per replicate, compared using a two-tailed t-test, all differences ns.

## Acknowledgements

We thank Vinay Godena, Caspar Baldwin, Alvaro Sanchez-Martinez, Tom Gleeson, Juliette Lee and Wing-Hei Au for help with various aspects of the fly work. Ana Terriente-Felix for help with stock maintenance and generating lines. Luis Gracia for developing the lifespan data analysis tool (https://flies.shinyapps.io/Rflies/). David Pate and Steve Drinkwater all their help setting up our fly lab. Carlo Viscomi and Hiran Prag for useful critical discussion. Richard Mann and Sumaira Zamurrad for the help with completing the fly work. Cristiane Benincà, Jordan Morris and the Wohl Cellular Imaging Centre at King’s College London for their help with microscopy. Karin H. Muller, Lyn Carter and Filomena Gallo for their help with TEM at the Cambridge Advanced Imaging Center (CAIC), and Jane Stinchcombe and Sam Loh for help interpreting this data. We thank Megan Oliva and Erin Barnhart and her lab for their critical reading of the manuscript. We thank Isabel Palacios for the Aβ-Arctic flies and Luca Scorrano and Valentina Debattisti for the synthetic tether line. Other stocks were obtained from the Bloomington Drosophila Stock Center which is supported by grant NIH P40OD018537.

## Author Contributions

VLH, LM-F, SA, JP, FM and AV designed and/or performed experiments, and analysed the data. VLH wrote the manuscript with input from all authors. AJW, AV and FP supervised the work.

